# Single-cell triple-omics uncovers DNA methylation as key feature of stemness in the healthy and ischemic adult brain

**DOI:** 10.1101/2022.07.13.499860

**Authors:** Lukas PM Kremer, Santiago Cerrizuela, Mohammad Eid Al Shukairi, Tobias Ellinger, Jannes Straub, Sascha Dehler, Aylin Korkmaz, Dieter Weichenhan, Christoph Plass, Simon Anders, Ana Martin-Villalba

## Abstract

Stem cells in the adult brain are specialized astrocytes capable of generating neurons and glial cells. While neural stem cells (NSCs) and common astrocytes have clearly distinct functions, they share highly similar transcriptome profiles. How stemness is molecularly encoded is therefore unclear. Here we use single-cell NMT-seq to simultaneously characterize the transcriptome, DNA methylome and chromatin accessibility of astrocytes and the NSC lineage in the healthy and ischemic brain. Our data reveal distinct methylation profiles associated with either astrocyte or stem cell function. Stemness is conferred by methylation of astrocyte genes and demethylation of neurogenic genes that are expressed only later. Surprisingly, ischemic injury unlocks the stemness-methylome in common astrocytes enabling generation of neuroblasts. Furthermore, we show that oligodendrocytes employ Tet-mediated demethylation to regulate expression of myelin-related genes, many of which are abnormally methylated in multiple sclerosis. Overall, we show that DNA methylation is a promising target for regenerative medicine.

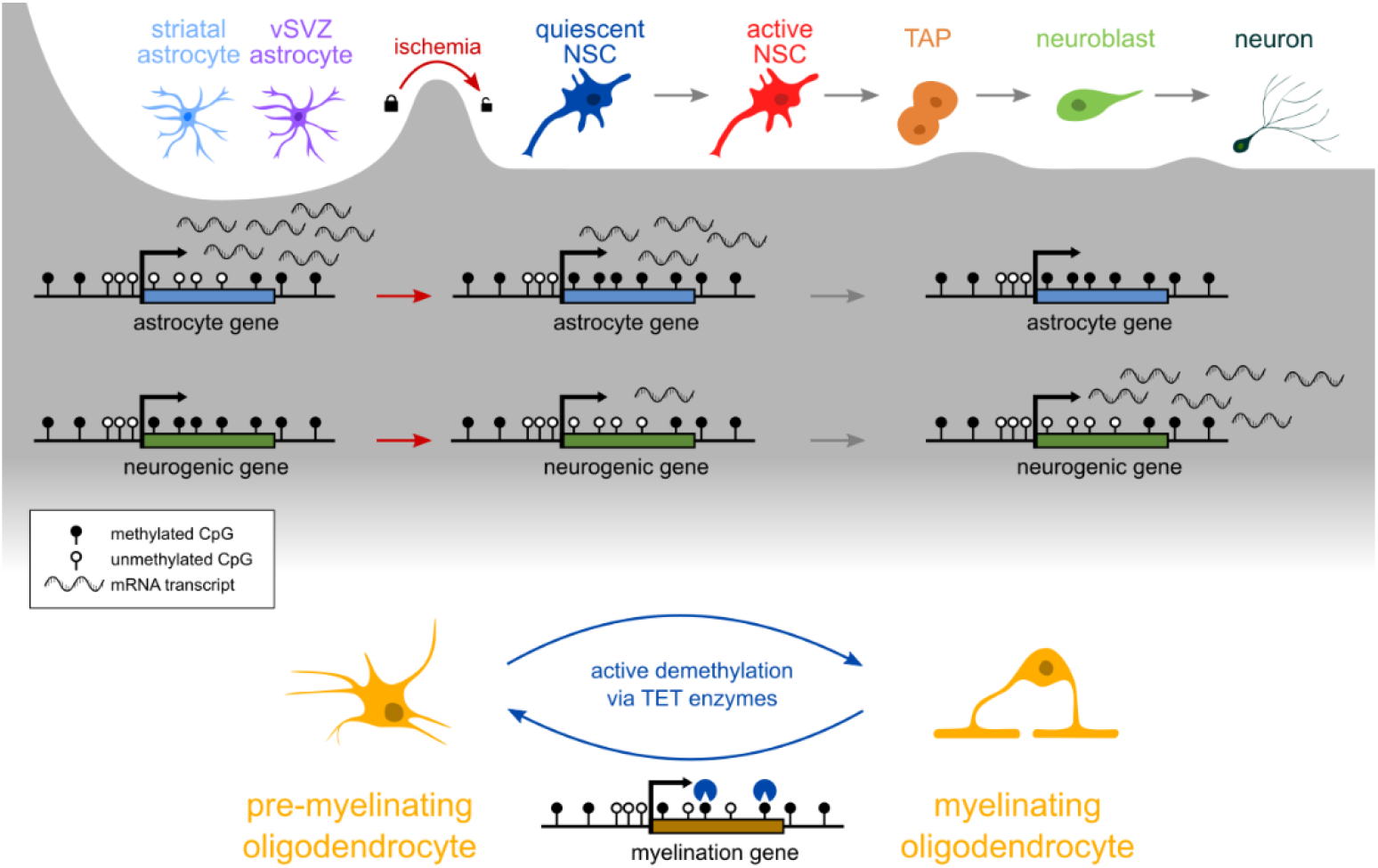

## Introduction

It was long thought that mammalian brains lose the ability to generate new neurons upon adulthood. Today we know that adult neurogenesis occurs but is limited to specialized niches including the dentate gyrus and the ventricular-subventricular zone (vSVZ). In the murine vSVZ, specialized astrocytes act as adult neural stem cells (NSCs) that reside in the wall of the lateral ventricles (Fig 1a). Upon activation, NSCs become transit-amplifying progenitors (TAPs) that undergo multiple rounds of division and give rise to neuroblasts (NBs). These then migrate along the rostral migratory stream (RMS) to the olfactory bulb (OB), where they differentiate into interneurons that integrate into the existing neural circuitry. Albeit to a much lesser extent, NSCs also give rise to glia, including other types of astrocytes and oligodendrocytes (Delgado et al., 2021; Lim & Alvarez-Buylla, 2016; Sohn et al., 2015).

**Figure 1.**
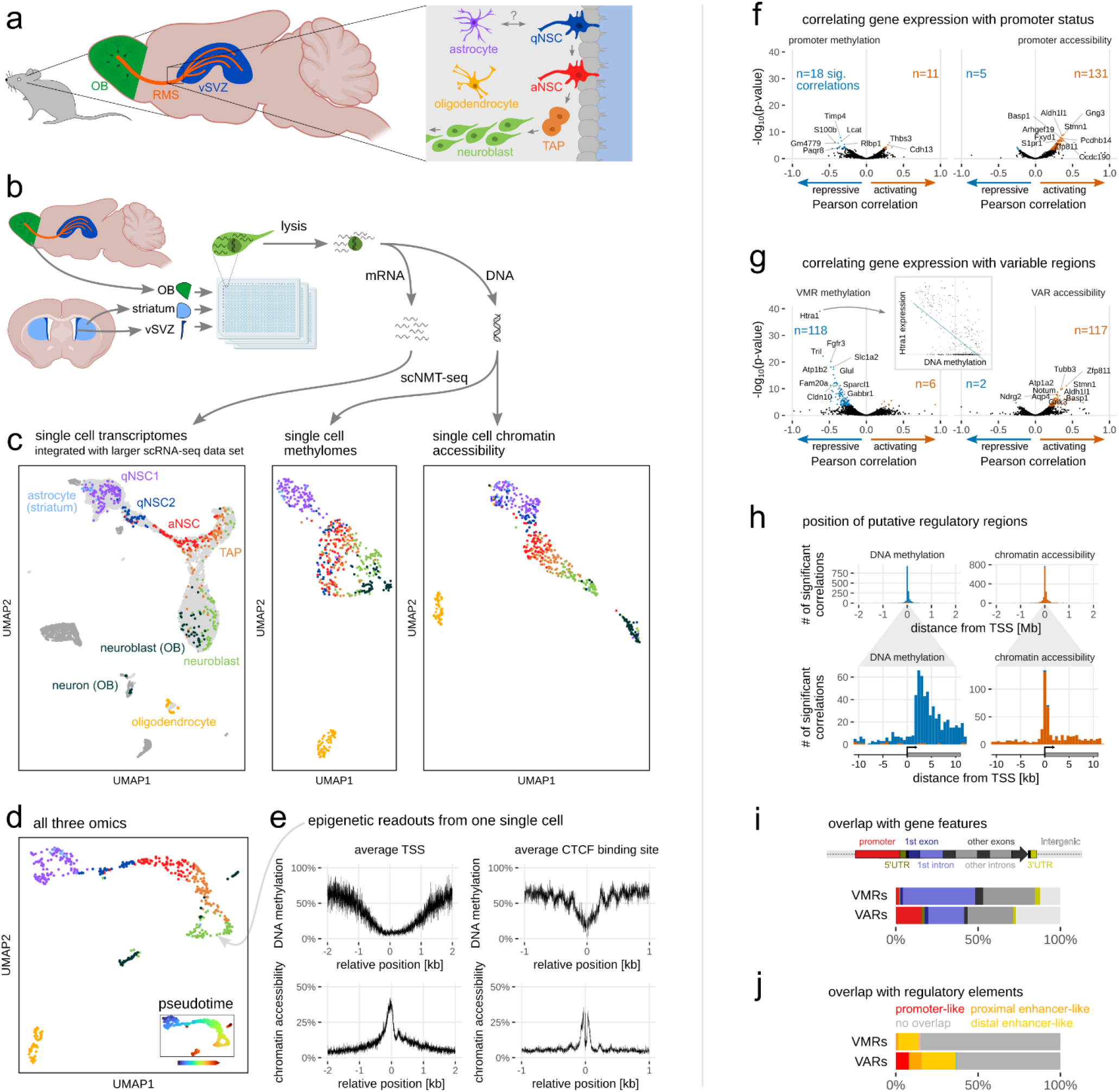
Single-cell triple-omics of the adult neural stem cell lineage. (**a**) The ventricular-subventricular zone (vSVZ) is populated by several cells including oligodendrocytes and two types of astrocytes, namely parenchymal astrocytes and neural stem cells. Quiescent neural stem cells (qNSC) may be activated (aNSC) to become rapidly dividing transit-amplifying progenitors (TAPs). TAPs then give rise to neuroblasts which migrate via the rostral migratory stream (RMS) to the olfactory bulb (OB), where they differentiate into neurons. (**b**) Schematic of the workflow to obtain scNMT-seq data from three brain regions. (**c**) UMAPs of each molecular layer, color coded as in *a*. For the transcriptomics data, the cells were integrated with the larger data set of Carvajal-Ibañez et al. (2022), which consists of cells from the vSVZ (light gray) and OB (dark gray). (**d**) UMAP and pseudotime assignments based on data from all three molecular layers. (**e**) Average methylation and chromatin accessibility levels around transcription start sites (TSS) and CTCF-binding sites of a single neuroblast. (**f**) Correlation of promoter methylation (left) and promoter accessibility (right) with gene expression. Negative correlations (blue) indicate a repressive effect of methylation/accessibility while positive correlations (orange) indicate activation. n: number of significant correlations after Benjamini-Hochberg adjustment. (**g**) Correlation of VMR (variably methylated region) methylation and VAR (variably accessible region) accessibility with expression of the nearest gene. (**h**) Distance histogram of all significant correlations between gene expression and all VMRs and VARs within 2 Mb of the TSS. Negative correlations are orange, positive correlations are blue. The bottom panel is a zoom-in of the top panel. (**i, j**) Overlap of significantly correlating VMRs and VARs with gene features (**i**) and candidate cis-regulatory elements from ENCODE (**j**).

The advent of single-cell RNA-sequencing (scRNA-seq) enabled the characterization of gene expression changes along the neurogenic lineage at unprecedented resolution (Cebrian-Silla et al., 2021; Llorens-Bobadilla et al., 2015; Zywitza et al., 2018). These studies showed that NSCs can be found in a quiescent or an active state. In the quiescent state, NSCs express genes associated with their astrocyte phenotype, including genes involved in lipid metabolism and glycolysis, which are gradually downregulated during the transition into the active state. Thus, NSCs are transcriptionally very similar to other astrocytes including parenchymal astrocytes of the neighboring striatum and cortex, which are generally viewed as non-neurogenic. This observation raises the question of the importance of the transcriptional profile for the potential to generate progeny. At the same time, it raises hopes for regenerative medicine, which aims to recruit these astrocytes to replace lost neurons in different brain regions. Indeed, several recent in vivo studies have reported astrocyte-to-neuron conversion by ablation or overexpression of key factors in the hippocampus, cortex and striatum (Magnusson et al., 2020; Mattugini et al., 2019; Nato et al., 2015; Qian et al., 2020). Other studies report that injury alone is sufficient to induce neurogenesis in some striatal astrocytes (Duan et al., 2015; Magnusson et al., 2014), raising the question of whether all astrocytes have latent neurogenic potential that is merely blocked in homeostasis.

Recently, scRNA-seq methods were extended to allow simultaneous profiling of gene expression and epigenetic marks. Here we employ and further develop the recently reported single-cell nucleosome, methylome and transcriptome sequencing (scNMT-seq (Clark et al., 2018)), which characterizes, simultaneously for each cell, three modalities: transcriptome, chromatin accessibility, and DNA methylation. This allowed us to assess whether gene expression changes in the NSC lineage are underpinned by epigenetic changes. Furthermore, we compared NSCs (neurogenic vSVZ astrocytes) with non-neurogenic astrocytes from the neighboring striatum to determine whether NSC stemness is encoded in the epigenome.

### Single-cell triple-omics of the adult NSC lineage

We first examined the transcriptome, methylome and chromatin accessibility of NSCs and their progeny. To this end, we isolated NSCs and their progeny from the vSVZ via flow cytometry as previously described (Kalamakis et al., 2019). Furthermore, we isolated astrocytes (GLAST^+^) from the striatum, as well as neuroblasts (PSA-NCAM^high^) and neurons (PSA-NCAM^low^) from the late RMS and OB. These cells were subjected to scNMT-seq to quantify gene expression, genome-wide DNA methylation and genome-wide chromatin accessibility at single-cell resolution (Fig. 1b). We increased the throughput of the original scNMT-seq protocol from 96 cells to 384 cells per run and reduced the volume in order to reduce costs. Furthermore, we updated the scRNA-seq portion of the protocol from Smart-seq2 to the recently developed Smart-seq3 (Hagemann-Jensen et al., 2020). After stringent quality filtering, we obtained a total of 542 triple-omic cells with an average of 5,811 expressed genes and 678,186 observed genomic CpG dinucleotides per cell (Supplementary Fig. 1a, Supplementary Tables 1-2).

Integration of our single-cell transcriptomes with a larger scRNA-seq reference dataset of the vSVZ, striatum and OB (Carvajal-Ibañez et al., 2022) revealed a continuum comprising two previously described (Kalamakis et al., 2019) sub-groups of quiescent NSCs (qNSC1, qNSC2), active NSCs (aNSC), TAPs, neuroblasts and neurons, as well as several oligodendrocytes (Fig. 1c, left). Of note, data from each epigenomic molecular layer alone was also sufficient to distinguish these cell states (Fig. 1c), suggesting that both, DNA methylation and chromatin accessibility, exhibit dynamic changes along this lineage. We then used MOFA+ (Argelaguet et al., 2020), a statistical framework for integration of multi-omic single-cell data, to reduce the data to 15 dimensions that incorporate information from all three molecular layers. We used this representation of the data to compute a 2D embedding (uniform manifold approximation and projection, UMAP, Fig. 1d) and to order the cells according to their progression in the NSC lineage (“pseudotime”). Our cell state assignments and the pseudotime ordering agree with the definitions from the literature as indicated by the expression of common marker genes and known lineage transcription factors (Supplementary Fig. 1b). To assess the quality of our epigenomic data, we next quantified DNA methylation and chromatin accessibility at transcription start sites (TSS) and CTCF binding sites (Fig. 1e). As was previously reported, the average TSS is lowly methylated and accessible (Clark et al., 2018). The average CTCF binding site shows a similar pattern but has more pronounced nucleosome marks (Teif et al., 2014) and decreased accessibility where CTCF is bound.

### Gene expression is regulated by promoter accessibility and DNA methylation of the first intron

Next, we set out to identify genes whose expression is regulated by epigenetic features. To do so, we made use of the single-cell resolution of our data and correlated gene expression with promoter methylation, and promoter accessibility, for every gene (Fig. 1f). As expected, we found that promoter methylation is typically repressive (negative correlations) while the opposite is true for promoter accessibility. We found far more significant associations between promoter accessibility and gene expression, indicating that few genes are regulated by promoter methylation. A likely explanation is that most promoters are unmethylated regardless of gene expression (Fig. 1e). To search the whole genome for putative regulatory regions, we used a sliding window approach (Kremer et al., 2022) to identify variably methylated regions (VMRs) and variably accessible regions (VARs). We found that the nearest VMR is a far better predictor of gene expression than promoters (Fig. 1g). Most VMRs occur inside the gene body about 3 kb downstream of the TSS (Fig. 1h) in the first intron (Fig. 1i). Only few VMRs overlap with known candidate cis-regulatory elements from ENCODE (Moore et al., 2020) (Fig. 1j).

**Supplementary Figure 1.**
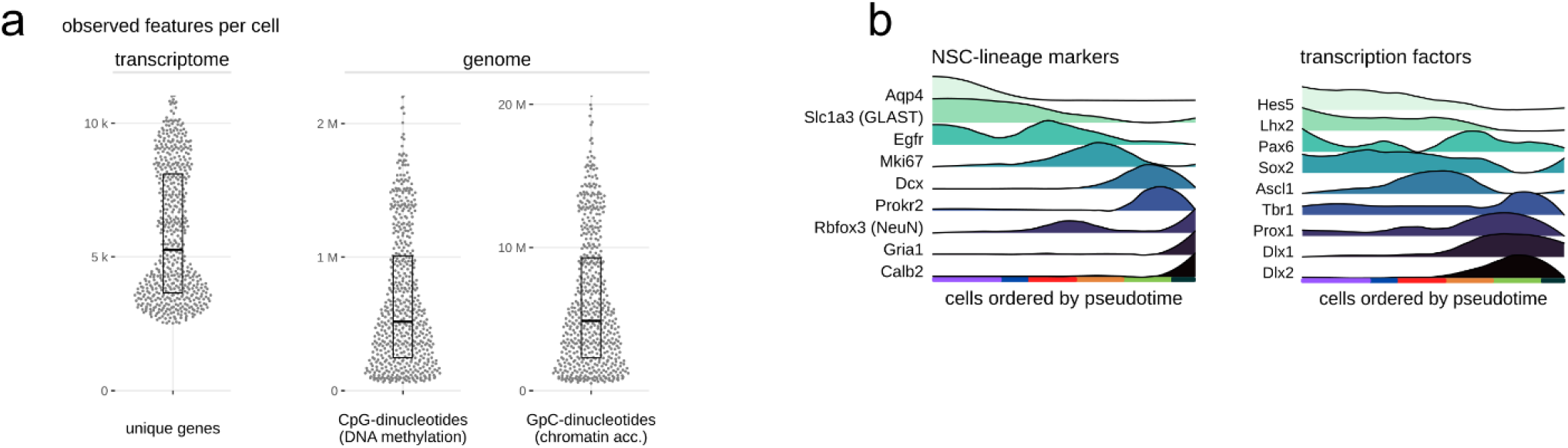
Quality measures and marker expression. (**a**) scNMT-seq quality metrics for all cells that passed quality filtering. “Unique genes” refers to the number of genes with at least one sequencing read per cell. “CpG/GpC-dinucleotides” refers to the number of methylation sites with sequencing coverage and thus known methylation status. (**b**) Gene expression of key marker genes and transcription factors along pseudotime.

### Adult NSC differentiation includes two waves of DNA methylation change

Several scRNA-seq studies (Cebrian-Silla et al., 2021; Llorens-Bobadilla et al., 2015; Zywitza et al., 2018) demonstrated that NSC differentiation is characterized by a series of gene expression changes, but whether this entails changes in DNA methylation has yet to be determined. To quantify the pace of changes along the lineage, we binned cells in pseudotime and calculated, for each of the three modalities, the correlation between the omics profiles of the pseudotime bins (Fig. 2a-b, Supplementary Fig. 2a). In these correlation heatmaps, large bright squares indicate groups of similar cells, and thus pseudo-time ranges with little change. Breaks between these squares indicate strong rapid change. As expected, the rapid changes in the transcriptomic profile coincide with the activation of NSCs (qNSC to aNSC) and with the differentiation of TAPs to NBs. The methylation heatmap, in contrast, shows a markedly different pattern, characterized by a very clear separation of qNSC1 and qNSC2 that was neither apparent in gene expression nor chromatin accessibility. Remarkably, qNSC1 cells shared the same methylome with striatal astrocytes, and will therefore henceforth be referred to as vSVZ astrocytes. The heatmap also showed several methylation changes in late TAPs and early neurons.

**Figure 2.**
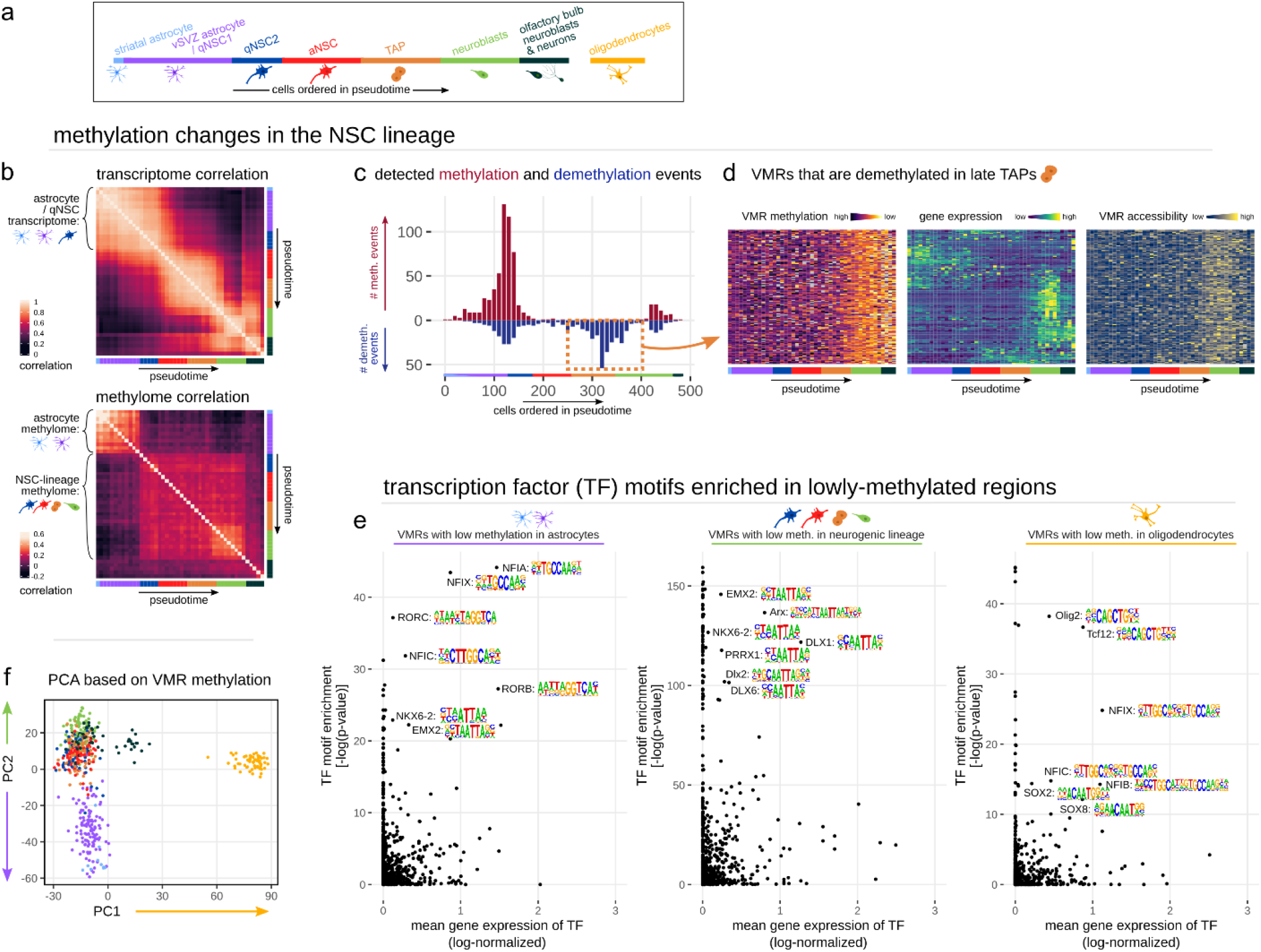
Adult neurogenesis involves a major and a minor wave of methylation change. (**a**) Schematic explaining the color code and pseudotime ordering of cells. (**b**) Correlation matrix of gene expression and DNA methylation. Cells are ordered and binned according to pseudotime, with each bin containing 10 cells on average. (**c**) Histogram of inferred pseudotime points at which VMRs become methylated (red) or demethylated (blue). (**d**) Expression and epigenetic state of genes intersecting with VMRs that become demethylated in late TAPs (dashed orange box in *c*). Rows are ordered by hierarchical clustering on gene expression values. Note that most of these genes are expressed in neuroblasts. (**e, f**) Motif enrichment of VMRs with low methylation in astrocytes, neuroblasts or oligodendrocytes as identified by PCA. (**e**) Scatterplots of transcription factors, showing the TF motif’s enrichment p-values on the y axis, and the TF mean gene expression in the respective cell population on the x axis. (**f**) PCA of single-cell methylomes. PC1 and PC2 separate oligodendrocytes and astrocytes, respectively, from the other cells, and were thus used to identify the regions used for motif enrichment in *e*.

Fitting a step function to the methylation values of each VMR across pseudotime revealed a first major wave of both methylation and demethylation in the transition from astrocyte to qNSC2, and a second wave of demethylation in late TAPs (Fig. 2c). A closer look at genes affected by demethylation in this second wave indicates that they are predominantly expressed in neuroblasts (Fig. 2d), suggesting that demethylation in late TAPs licenses neuroblast genes for later expression. Curiously, demethylation of these regions is accompanied by an only transient period of chromatin accessibility. In most cases, accessibility coincides with gene expression, while low methylation persists even in those genes that are downregulated at the neuron stage (Fig. 2d).

To observe general patterns in the regulation of other state-specific genes, we used differential gene expression analysis to determine 100 marker genes for each stage of the NSC lineage (Supplementary Table 3). We then visualized their mean expression, as well as the mean methylation and accessibility of nearby VMRs and promoters (Supplementary Fig. 2b). We observe three clear patterns: First, while gene expression of astrocyte markers fades gradually, the methylation of nearby VMRs is clearly distinct between vSVZ astrocytes and qNSC2. Second, the promoters of TAP markers, many of which are involved in mitosis, are demethylated and accessible in all cell states that we studied. Third, we observed that oligodendrocyte marker expression is clearly underpinned by low methylation and increased accessibility of both promoters and VMRs. Interestingly, accessibility of oligodendrocyte markers shows a slight increase upon NSC activation, which might be due to the presence of oligodendrocyte-fated cells in the lineage. Markers for intermediate cell states, however, show weaker, more ambiguous epigenetic patterns, suggesting that regulation by accessibility or methylation is less important for these genes.

Finally, we asked whether the observed methylation differences are the result of transcription factor (TF)-guided (de)methylation as previously described (Donaghey et al., 2018; Reizel et al., 2021; Suzuki et al., 2017). To address this question, we used TF motif enrichment on regions that are demethylated in either oligodendrocytes, astrocytes, or the neurogenic lineage as determined by PCA (Fig. 2e-f). To narrow the list of candidate TFs, we furthermore considered the average gene expression of each TF in the respective cell population. Regions demethylated in astrocytes frequently contain the motif of one or more nuclear factors. Among them is NFIA, which is known to induce demethylation of the astrocyte marker GFAP and is used to convert human induced pluripotent stem cell (iPSC)-derived NSCs to astrocytes (Tchieu et al., 2019), and NFIX, a TF that regulates NSC quiescence and suppresses oligodendrogenesis (B. Zhou et al., 2015). Regions lowly demethylated in oligodendrocytes were enriched for the motif of Olig2, a well-studied master regulator of oligodendrocyte cell identity (K. Zhang et al., 2022; Q. Zhou et al., 2000), as well as Tcf12, which may be involved in the known generation of oligodendrocyte-fated NSCs by Wnt ligands (Ortega et al., 2013).

**Supplementary Figure 2.**
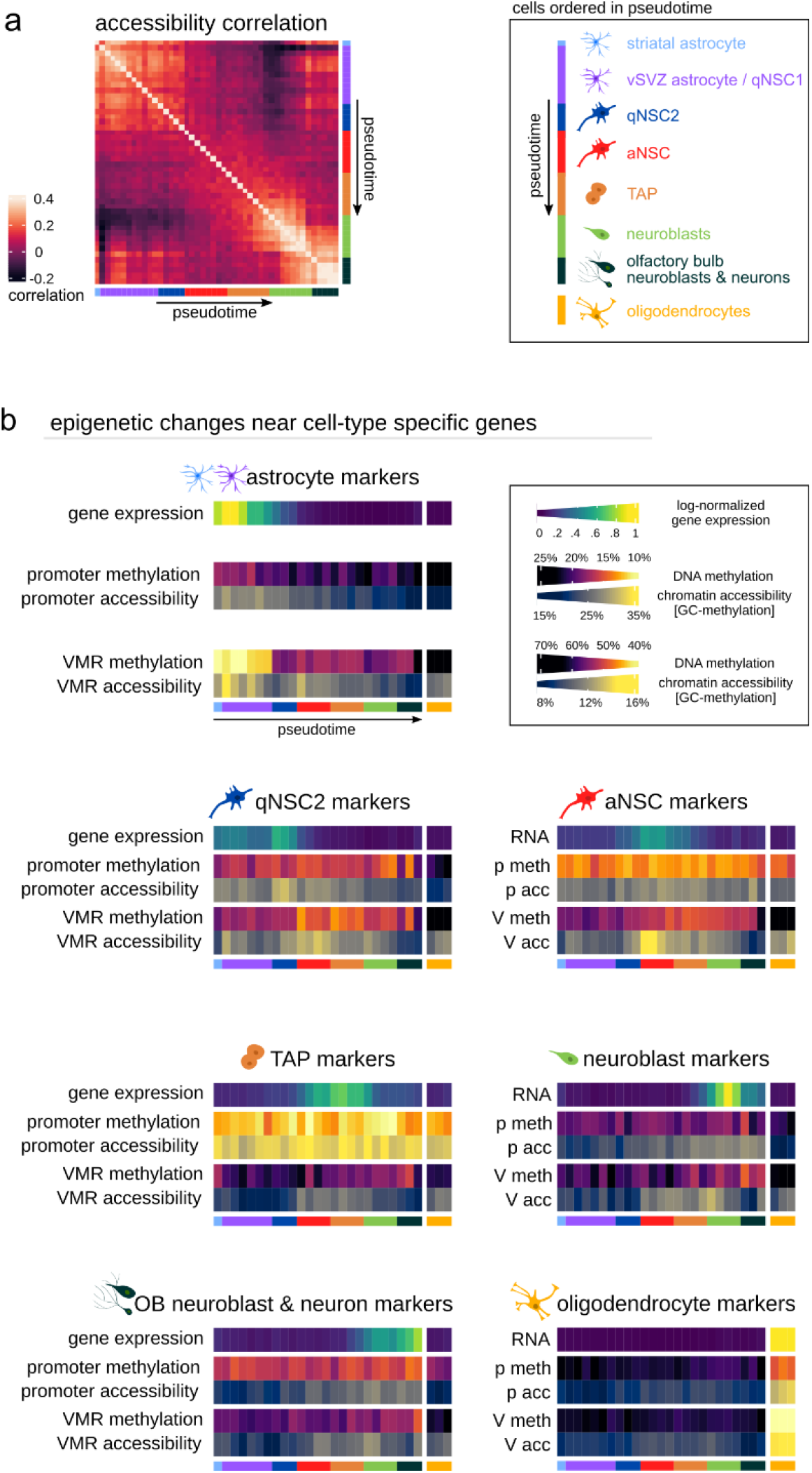
Epigenetic changes in pseudotime. (a) Correlation matrix of chromatin accessibility. Cells are ordered and binned according to pseudotime, with each bin containing 10 cells on average. (**b**) Expression and epigenetic status of genes expressed at different states of the NSC lineage. The top heatmap row indicates the average expression of 100 marker genes identified in a differential expression analysis of the single-cell transcriptomes. Middle and bottom rows indicate the average DNA methylation (meth) and chromatin accessibility (acc) at the 100 promoters (p) of these markers, and at 100 VMRs (V) overlapping marker gene bodies.

### Despite similar gene expression, DNA methylation strongly differs between NSCs and other astrocytes

We then focused on the substantial DNA methylation differences between astrocytes and qNSC2 (Fig. 2b,c,f). DMR detection identified lowly methylated regions (LMRs) in both astrocytes and qNSC2, and nearby genes tended to be more highly expressed in the respective subtype (Fig. 3a, Supplementary Table 4). Regions lowly methylated in astrocytes occur near genes involved in ion transport (Slca41a2), amino acid transport and metabolism (Slca1a2, Glul) and cholesterol metabolism (Lcat), among others (Fig. 3b). Since these GO terms represent fundamental astrocyte functions and since the corresponding regions are also lowly methylated in striatal astrocytes (Fig. 3c), we conclude that these regions are associated with an astrocyte cell identity and labeled them astrocyte LMRs.

**Figure 3.**
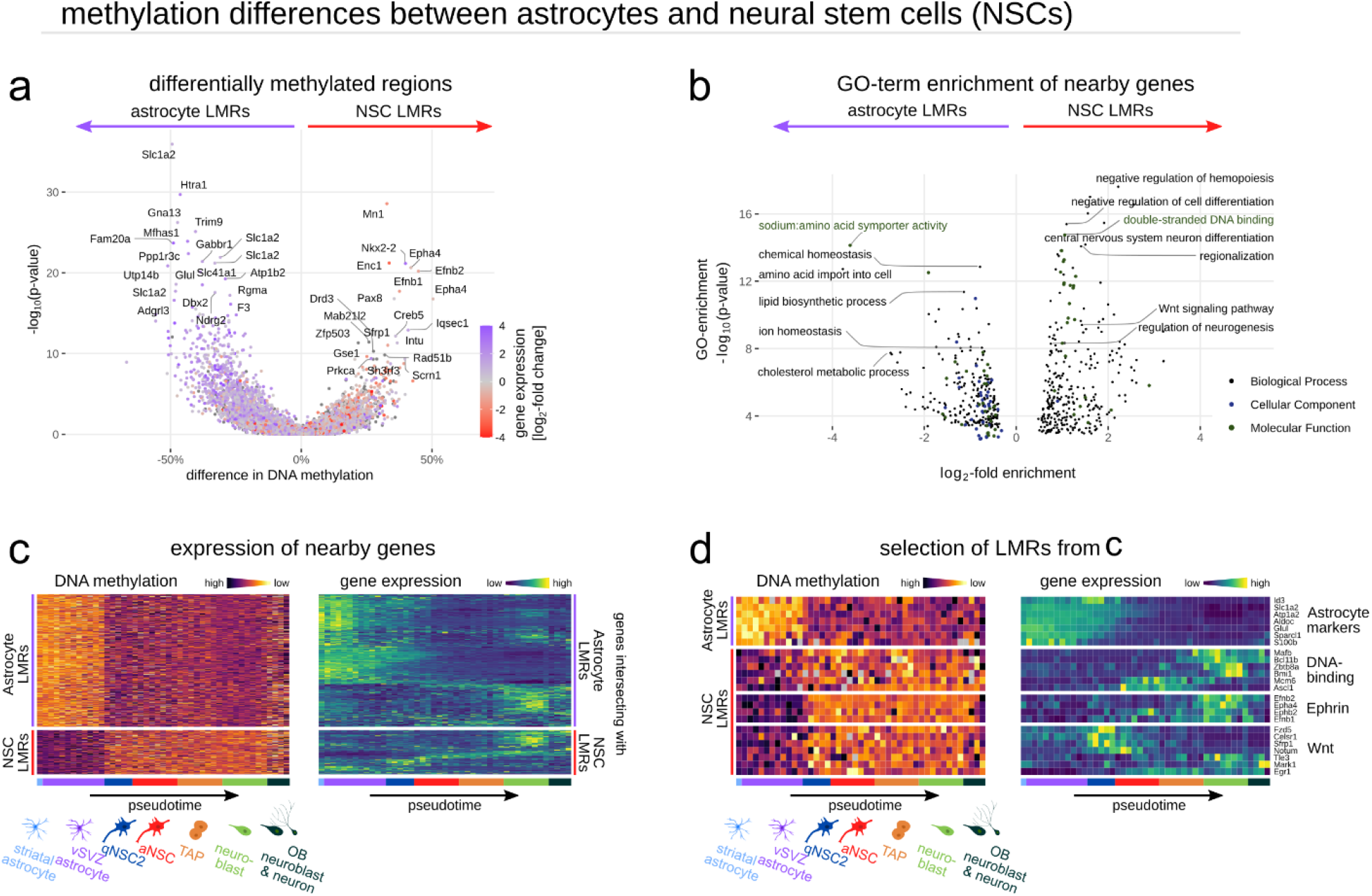
NSCs possess a pro-neurogenic methylome that clearly distinguishes them from common astrocytes. (**a**) Volcano plot of VMRs, tested for differential methylation between vSVZ astrocytes and NSCs. LMR: lowly-methylated region. (**b**) GO-term enrichment of genes near astrocyte LMRs and NSC LMRs identified in *a*. (**c**) Heatmap of LMR methylation (left) and expression of the nearest gene (right) along pseudotime. Rows are ordered by hierarchical clustering on gene expression values. (**d**) Selection of LMRs and genes from *c*. Note the clear separation of NSCs and astrocytes in the methylation data.

In contrast, genes near regions that are lowly methylated in qNSC2 are enriched for regulators of cell differentiation (e.g., Ascl1, Dlx1) and DNA binding (e.g., Nkx2-2, Pax8) (Fig. 3a-b). Among the top hits are also several genes of the ephrin signaling pathway (e.g., Efnb2, Epha4, Efnb1) that controls neuroblast migration and NSC quiescence (Holmberg et al., 2005; Jiao et al., 2008; Lim & Alvarez-Buylla, 2016; Nomura et al., 2010; Ricard et al., 2006). This implies that demethylation at these regions contributes to stem cell function. Hence, we labeled them NSC LMRs. We conclude that astrocytes and qNSC2 cells, two populations that are notoriously hard to distinguish by their transcriptome (Lim & Alvarez-Buylla, 2016), exhibit very distinct methylomes, tied to either astrocyte function in the case of vSVZ and striatal astrocytes, or to stem cell function in the case of qNSC2. This clear divide by their methylomes does not translate to the transcriptome since many astrocyte markers are expressed in both populations. Similarly, expression of genes near NSC LMRs also reaches maximum values at various points of the NSC lineage even though methylation is constantly low (Fig. 3c). This suggests that genes required at later states are epigenetically licensed for later expression already in NSCs, despite not yet being expressed. Among these genes are ephrin signaling genes expressed in neuroblasts, DNA-binding genes expressed after NSC activation and Wnt signaling genes of which some are specifically expressed in quiescent NSCs (Fig. 3d). To summarize, we found vast methylation differences between NSCs and other astrocytes, despite similar transcriptomes.

### Ischemic injury induces an NSC-like methylome in striatal astrocytes

We have previously shown that such an ischemic injury activates the exit of a quiescent state into an activation state, as assessed based on their transcriptomes (Llorens-Bobadilla et al., 2015). Only now, with the ability to differentiate vSVZ astrocytes from qNSC2, we can address if ischemia activates only qNSC2, or vSVZ astrocytes as well. Activation of astrocytes by an injury has been previously shown for striatal astrocytes (Magnusson et al., 2020), however, whether this involves full conversion of the astrocyte-methylome into the stemness-methylome is unknown. To test this, we challenged these cells by inducing transient global brain ischemia in 2-month-old mice. From these mice, we isolated GLAST^+^ astrocytes and NSCs from the vSVZ and striatum for scNMT-seq, yielding another 614 triple-omic cells (Fig. 4a, Supplementary Table 1). To ensure that the cells isolated from the striatum had not migrated from the vSVZ, we induced YFP expression exclusively in vSVZ NSCs. To this end, we used a transgenic mouse line expressing Cre/ERT2 recombinase under the promoter of Nr2e1 (Tlx), a marker of vSVZ NSCs (Liu et al., 2008). Upon tamoxifen injection 2 weeks before ischemia, the Cre/ERT2 recombinase is induced in Tlx^+^ cells, which leads to excision of a STOP cassette upstream of YFP. Reassuringly, we detected YFP^+^GLAST^+^-cells only in the vSVZ and not in the striatum (Supplementary Fig. 3a-d).

**Figure 4.**
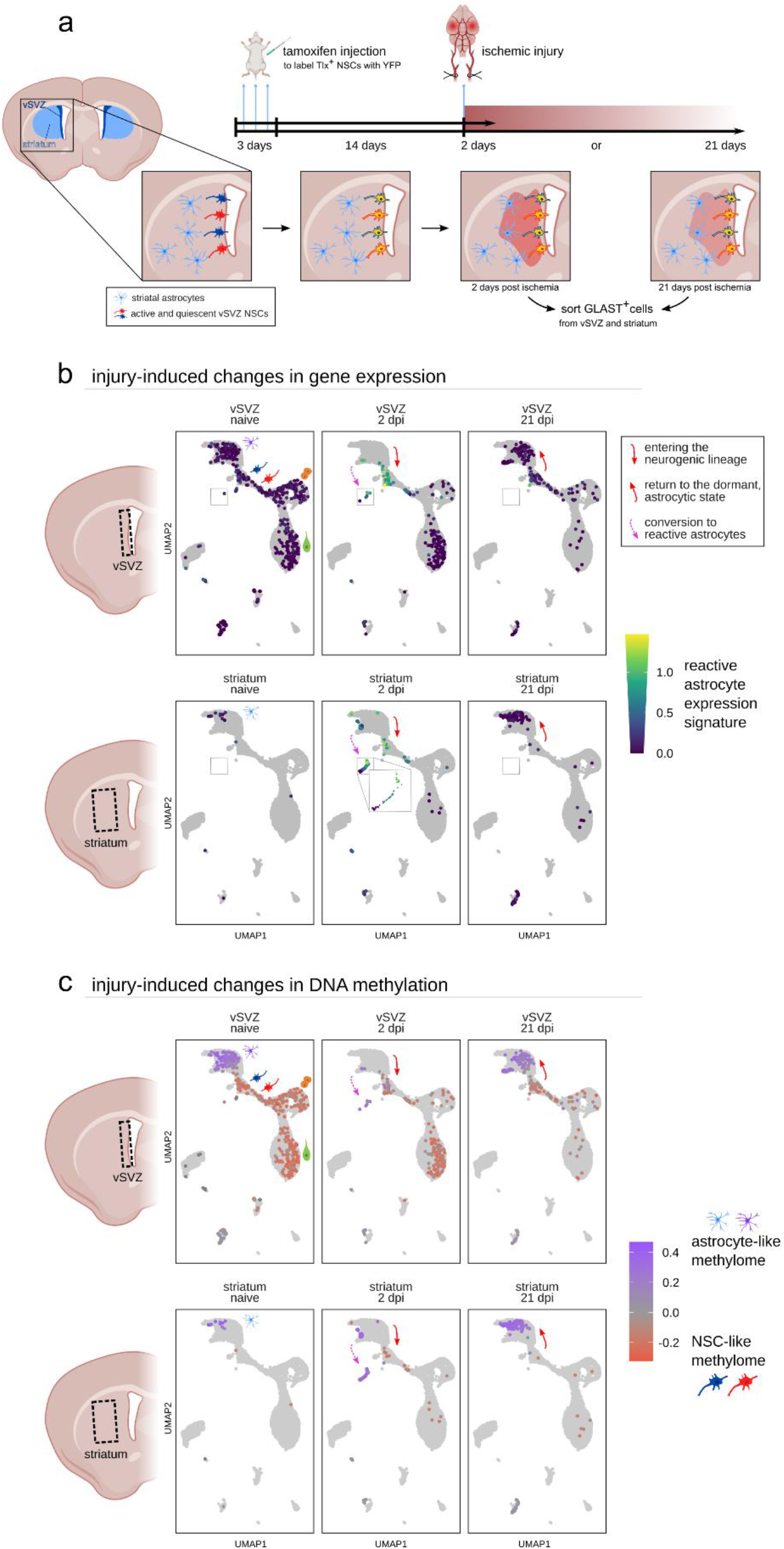
Ischemic injury induces an NSC-like methylome in striatal astrocytes. (**a**) Schematic of the experiment to assess the effects of ischemic injury on GLAST^+^ cells in the vSVZ (astrocytes and NSCs) and striatum (astrocytes). Both tissues were sequenced at two timepoints: 2 dpi and 21 dpi. TAM is injected to label Tlx^+^ NSCs via Cre-inducible YFP expression to detect potential NSCs that may migrate to the striatum. (**b**) Comparison of the cell states 2 dpi and 21 dpi with the cell states in naïve mice. Cells are colored by mean log-normalized expression of reactive astrocyte marker genes from Liddelow et al., 2017. (**c**) Same data, now colored according to summarized methylation of NSC LMRs and astrocyte LMRs from Fig. 3. The depicted score is the difference between the average methylation of all astrocyte LMRs and the average methylation of all NSC LMRs (also see Supplementary Fig. 4e-f).

Projecting the ischemic transcriptomes again onto the feature space of the naïve cells from Carvajal-Ibañez et al. (2022) confirmed an exit of vSVZ astrocytes into the qNSC2 state 2 days post ischemia (dpi) in the vSVZ (Llorens-Bobadilla et al., 2015) (Fig. 4b). The pool of vSVZ astrocytes was replenished 21 dpi, indicating a return to a dormant state. In the case of striatal astrocytes, the majority converted into the reactive astrocyte state, a state characterized by pronounced changes in function, morphology and gene expression (Anderson et al., 2016; Liddelow et al., 2017). However, some of them entered the neurogenic lineage, similarly to vSVZ astrocytes. Among reactive astrocytes, two subtypes have been previously described: A1 and A2 (Liddelow et al., 2017). Based on gene expression, we found that striatal astrocytes that enter the neurogenic lineage more closely resemble A1 astrocytes, while others rather exhibit an A2 profile (Fig. 4b, Supplementary Fig. 4a-d). Notably, the A1 profile has typically been considered neurotoxic (Liddelow et al., 2017). Our data indicate that directly after injury, A1 astrocytes feature a neurogenic/stemness state.

Most importantly, both these neurogenic striatal astrocytes and vSVZ astrocytes activated by ischemia exhibit an NSC-like methylome, characterized by high methylation of astrocyte LMRs and low methylation of NSC LMRs (Fig. 4c, Supplementary Fig. 4e-f). This finding indicates that ischemia not only causes activation of a neurogenic transcriptional program in astrocytes, but also substantial epigenetic remodeling including the demethylation of neuroblast-specific genes.

Furthermore, a direct methylome comparison of naïve astrocytes with post-ischemic astrocytes revealed demethylation near genes associated with the GO-term “ruffle membrane” (Supplementary Fig. 5a-c). This suggests a potential role for demethylation in reactive astrocytes, which are known to extend cellular processes with characteristic membrane ruffles into cell-free space at a wound edge (Schiweck et al., 2018). Curiously, 21 days after ischemia, the most significantly demethylated site is located in the gene body of Dnmt3a, the methyltransferase responsible for *de novo* DNA methylation (Supplementary Fig. 5d-f).

**Supplementary Figure 3.**
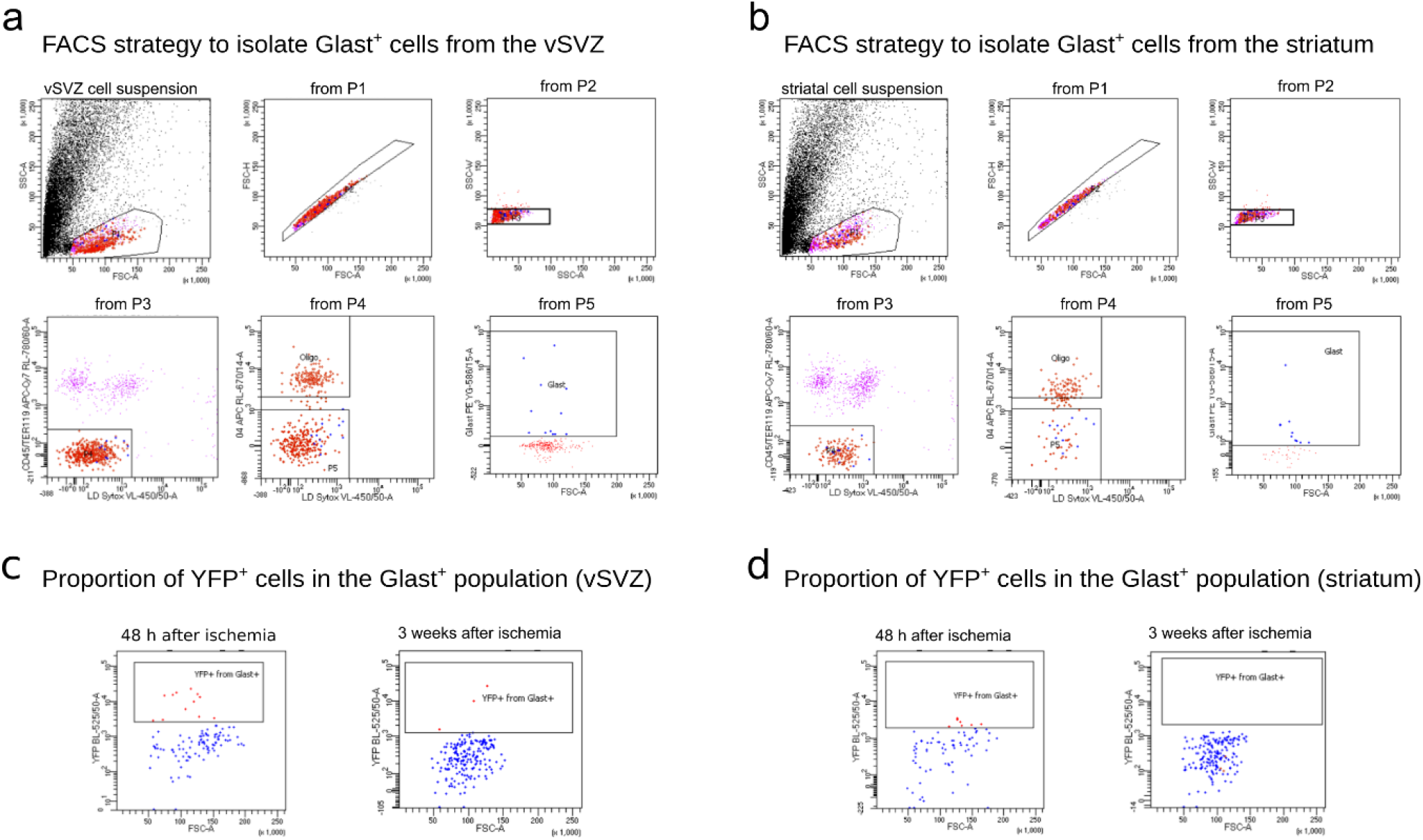
Cell sorting demonstrates that cells captured from the striatum did not migrate from the vSVZ. **(a**,**b)** Strategy for sorting Glast^+^ cells from the vSVZ (**a**) and striatum (**b**). Striatal Glast^+^ cells correspond to astrocytes, while vSVZ Glast^+^ cells correspond to NSCs and astrocytes. Dead cells and CD45/O4/Ter119 positive cells were excluded. **(c**,**d)** The YFP intensity from the Glast^+^ cells was also captured by index sorting. Dot plots show the fraction of YFP^+^ cells from the Glast^+^ population, being higher in the vSVZ sample 2 dpi. Axis information: Antigen names are written on each axis, followed by the filter used to measure the signal. FACS: Fluorescence-activated cell sorting. SSC-A: Side scatter area. FSC-A: forward scatter area. FSC-H: forward scatter height. RL-780/60-A: APC-Cy7 fluorochrome. VL-450/50-A: Pacific blue fluorochrome. RL-670/14-A: APC fluorochrome. YG-586/15-A: PE fluorochrome.

**Supplementary Figure 4.**
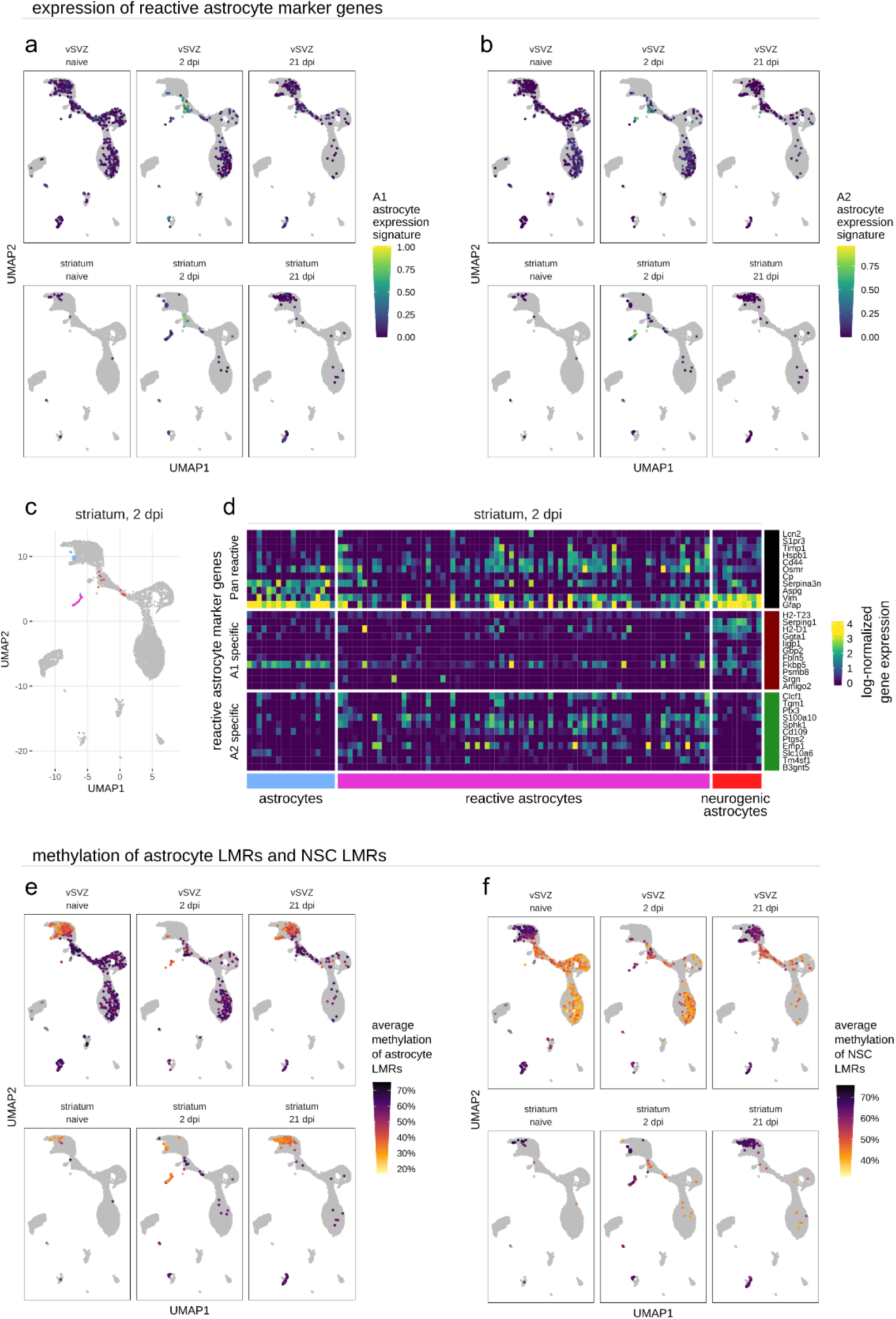
Expression of reactive astrocyte genes and methylation of NSC/astrocyte LMRs. (**a, b**) Mean log-normalized expression of marker genes for the two subtypes of reactive astrocytes (a: A1, b: A2) listed by Liddelow et al., 2017. (**c**) Clustering of striatal astrocytes 2 dpi. (**d**) Expression of reactive astrocyte marker genes. Columns correspond to the cells highlighted in *c*. (**e, f**) Average methylation of astrocyte LMRs (**e**) and NSC LMRs (**f**) from Fig. 3.

**Supplementary Figure 5.**
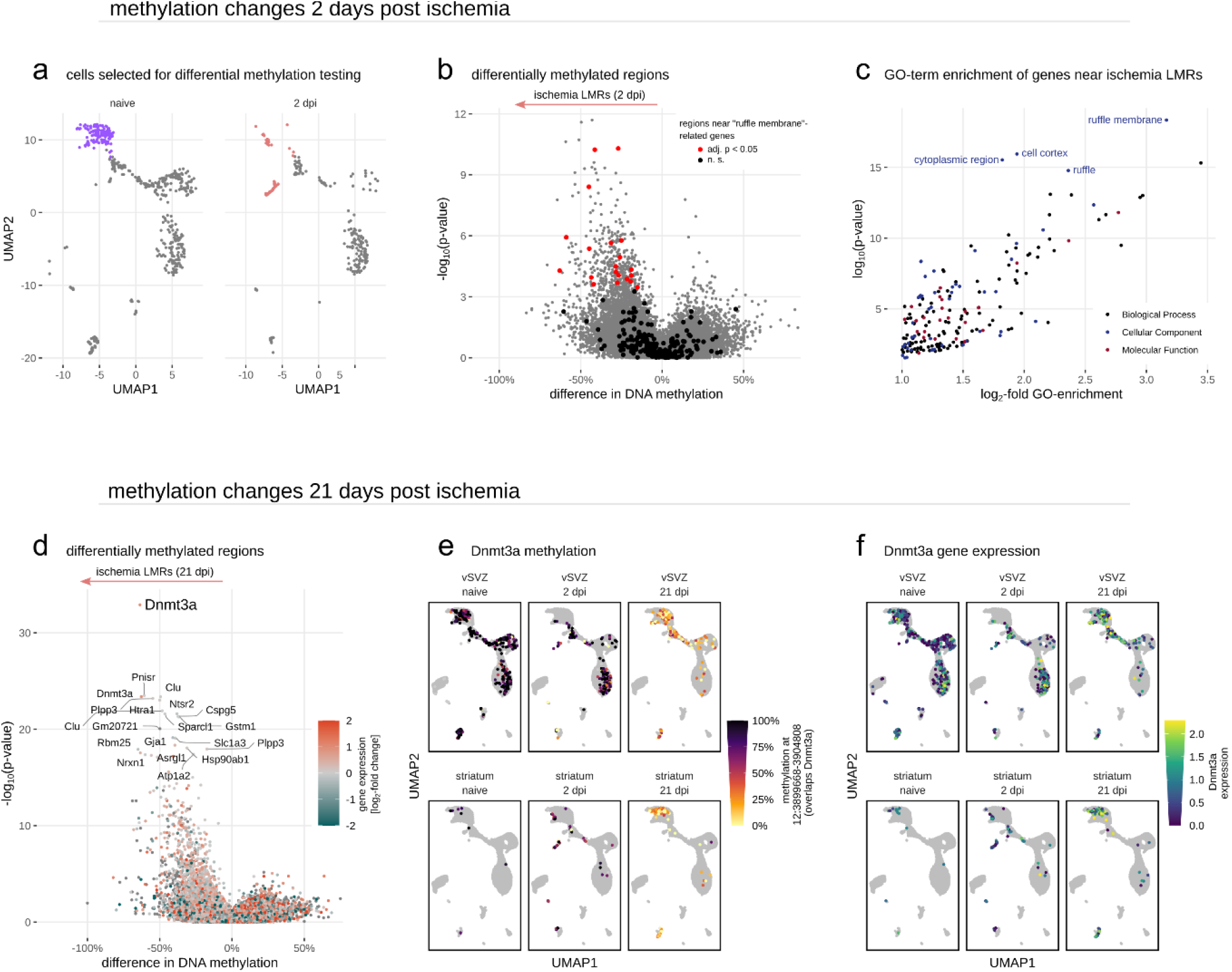
Ischemia induces demethylation near Dnmt3a and near genes involved in ruffle membranes. (**a-c**) Methylation changes observed in astrocytes 2 days post ischemia (dpi). (**a**) Naïve striatal astrocytes and vSVZ astrocytes (purple) were compared to the same cell populations 2 dpi (highlighted in red; also includes reactive astrocytes). (**b**) Volcano plot of VMRs, tested for differential methylation. (**c**) GO-term enrichment of genes near LMRs identified in *b*. (**d-f**) Methylation changes observed in astrocytes 21 dpi. (**d**) Volcano plot of VMRs, tested for differential methylation between naïve astrocytes (striatal astrocytes, vSVZ astrocytes) and astrocytes 21 dpi. (**e**) Methylation status of the genomic region 12:3899668-3904808, identified in *d*, that is located in the Dnmt3a gene body. (**f**) Log-normalized gene expression of Dnmt3a. Outlier values (top 2%) were pulled in.

### Oligodendrocytes employ TET enzymes to regulate the expression of myelin-related genes

In addition to LMRs identified in the neurogenic lineage, we also identified oligodendrocyte specific LMRs (Fig. 5a). Several of these regions are located close to commonly used oligodendrocyte markers (Mbp, Mag, Mobp, Sox10, …) involved in myelination, the primary function of oligodendrocytes (Fig. 5b). Next, we set out to address the molecular mechanism responsible for the methylation changes we identified in the NSC lineage and oligodendrocytes. DNA methylation can be removed either actively or passively. Passive DNA methylation occurs when the DNA methylation maintenance machinery is inhibited or absent during DNA replication. In contrast, active demethylation is a targeted, stepwise oxidation of 5-methylcytosines catalyzed by enzymes of the Ten-Eleven Translocation (TET) family (reviewed by Wu and Zhang, 2017) (Fig. 5c, top). Active demethylation by TET enzymes plays a pivotal role in animal development by controlling the expression of lineage-specific genes in embryonic stem cells and embryonic neural progenitors (Argelaguet et al., 2019; Bogdanović et al., 2016; Reizel et al., 2018). TET depletion experiments also demonstrated that TET enzymes affect adult NSCs in the dentate gyrus of the hippocampus (Guo et al., 2011; Li et al., 2017). Thus, we examined whether the LMRs we observed are the result of TET-mediated demethylation.

**Figure 5.**
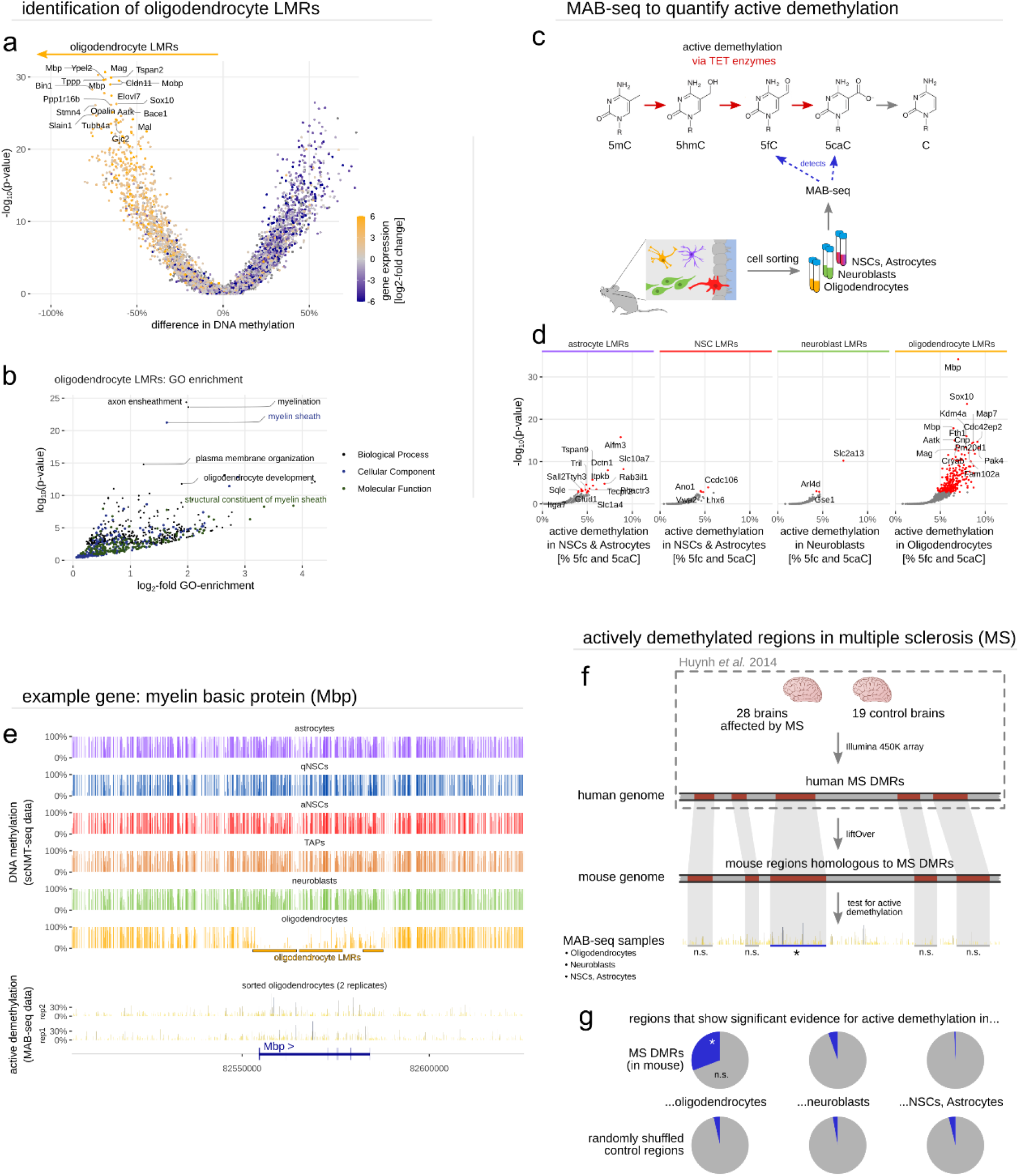
Oligodendrocytes employ TET enzymes to regulate expression of myelin-related genes. (**a**) Volcano plot of VMRs, tested for differential methylation between oligodendrocytes and all other cells. (**b**) GO-term enrichment of genes near oligodendrocyte LMRs identified in *a*. (**c**) Schematic of TET-mediated active demethylation (top) (5mC, 5-methylcytosine; 5hmc, 5-hydroxymethylcytosine; 5fC, 5-formylcytosine; 5caC, 5-carboxylcytosine) and our experimental workflow to detect active demethylation with MAB-seq (bottom). (**d**) Quantification of active demethylation in cell type-specific LMRs. The genome coordinates of LMRs were previously identified in scNMT-seq data. These coordinates are assessed in the MAB-seq data to quantify active demethylation at LMRs. (**e**) DNA methylation (top) and remnants of active demethylation (bottom) near myelin basic protein (Mbp). (**f**) Quantification of active demethylation in genomic regions orthologous to MS DMRs reported by Huynh et al., 2014, using our MAB-seq data. (**g**) 30.8% of MS DMRs show significant evidence for active demethylation in murine oligodendrocytes, in contrast to other cell types (top) or random control intervals of the same size distribution (bottom).

To test this hypothesis, we used methylase-assisted bisulfite sequencing (MAB-seq) (H. Wu et al., 2014). MAB-seq allows genome-wide detection of 5-formylcytosine (5fC) and 5-carboxylcytosine (5caC), two oxidized 5-methylcytosine (5mC) bases indicative of recent TET activity. We performed MAB-seq on three populations freshly isolated from the vSVZ: GLAST^+^CD9^+^ cells including vSVZ astrocytes and NSCs, PSA-NCAM^+^ neuroblasts, as well as Prom1^+^/Glast^-^ oligodendrocytes (Fig. 5c). We next used our MAB-seq data to test all LMRs for traces of active demethylation. Indeed, we detected 5fC and 5caC in several astrocyte LMRs, as well as in a small number of NSC LMRs and neuroblast LMRs (Fig. 5d). We conclude that some, but not all demethylation events in the NSC lineage are mediated by TET enzymes. To our surprise, however, we detected active demethylation in hundreds of oligodendrocyte LMRs. Nearby genes include Sox10, a transcription factor that acts as a master regulator of oligodendrocyte fate (Mokhtarzadeh Khanghahi et al., 2018; Pozniak et al., 2010; K. Zhang et al., 2022), and myelin basic protein (Mbp) (Fig. 5e), the second most abundant protein in the myelin sheath with important functions in myelin sheath formation and stabilization (Boggs, 2006). These intriguing results indicate that active demethylation might be involved in the regulation of myelin-related genes in adult brains.

Therefore, we wondered whether TET-mediated demethylation might be compromised in demyelinating diseases such as multiple sclerosis (MS). Huynh et al. (2014) previously showed that human brains affected by MS harbor regions with different methylation levels compared to healthy controls (MS DMRs). Notably, several mouse regions orthologous to human MS DMRs showed clear evidence of TET-mediated demethylation in our MAB-seq data (Fig. 5f-g). Altogether our data indicate that oligodendrocytes regulate myelination through active demethylation of regions ultimately regulating expression of myelinating factors, and that regulation of the methylation state of some of these regions is compromised in brains from MS patients.

## Discussion

### Insights into the role of DNA methylation in adult tissues

The past decades have demonstrated that new leaps in our understanding of DNA methylation are often driven by technological innovations such as bisulfite sequencing (Frommer et al., 1992; Mattei et al., 2022). In a similar manner, our *in vivo* assessment of DNA methylation in cell populations of the adult NSC lineage was only possible due to recent advances in single-cell sequencing and multi-omics. Both the single-cell resolution of the data, as well as the added transcriptomic layer, allowed us to quantify DNA methylation in precisely defined cell populations. Thus, with scNMT-seq, we have been able to free ourselves from the need for pure cell populations that rely on a large panel of excellent FACS surface markers, and from the need for cell culture systems known to alter DNA methylation (Antequera et al., 1990; Franzen et al., 2021). Nonetheless, single-cell multi-omics also introduced new challenges such as the sparsity of the data and the modest cell numbers that are currently achievable. To conduct this study, we tackled both problems by developing new computational methods for single-cell methylation data (Kremer et al., 2022) and miniaturizing the scNMT-seq protocol (Cerrizuela et al., 2022).

The methylation changes we observed rarely occurred in promoters but were enriched for gene bodies including the first intron. This observation agrees with the current view that promoter methylation is generally low and less dynamic than methylation in cis-regulatory elements (Mattei et al., 2022; Meissner et al., 2008; Stadler et al., 2011). On the other hand, relatively few VMRs were located in known cis-regulatory elements. There are several possibilities that may explain this discrepancy. First, some VMRs may be located in un-annotated regulatory elements. Second, some VMRs may be introduced to silence cryptic intragenic promoters when a gene is upregulated (Neri et al., 2017). This would also explain why methylation of some VMRs is positively correlated with gene expression (Fig. 1g, Fig. 2d, Fig. 3c). Third, we note that not all regulatory methylation changes occur inside cis-regulatory elements. For instance, distal methylation change can result in the formation of activating or repressive chromatin loops (Bell & Felsenfeld, 2000; Teif et al., 2014). Finally, our data supports the currently underappreciated notion that DNA methylation downstream of the promoter, in the first intron, silences gene expression (Anastasiadi et al., 2018; Schlosberg et al., 2017).

Classically, DNA methylation was viewed as a repressive epigenetic mark that remains static once established (Mattei et al., 2022). In recent years, however, both views were overturned by studies demonstrating that DNA methylation is dynamic in embryonic development (Argelaguet et al., 2019; Bogdanović et al., 2016; Reizel et al., 2018) and in stem cells differentiating *in vitro* (T. Gu et al., 2018; Xie et al., 2013). In this work we demonstrated that the same principle also applies to NSCs differentiating in adult brains. Even more surprisingly, our results suggest that methylation also changes in response to environmental stimuli such as ischemia, and is actively modified to regulate myelination in oligodendrocytes in adulthood. These findings indicate that DNA methylation is even more dynamic than previously thought and that it may play a role in a wide range of biological processes in adult tissues.

### Active demethylation in oligodendrocytes

Our results demonstrate that oligodendrocytes employ TET enzymes to modify DNA methylation near genes required for myelination in adulthood. This result was unexpected, since changes in DNA methylation are usually associated with development and cellular differentiation. However, during the course of this study, Zhang et al. (2021) used a very different set of methods to show that Tet1 is required for both myelination in development, as well as remyelination after injury in adulthood. Taken together, these results suggest a novel, unexpected role for active demethylation in dynamically fine-tuning the expression of myelin-related genes. Our study lays the foundation for this emerging research topic by unveiling the regions targeted by TET enzymes in oligodendrocytes. Along the same line, we also observed that induction of reactive astrocytes involves demethylation near genes involved in membrane ruffles (Supplementary Fig. 5a-c). These two findings pose the question whether there is a more general, previously unappreciated role for DNA methylation in the re-organization of glial membranes.

### Implications for adult NSC and astrocyte biology

First and foremost, we believe that we made a crucial step toward understanding why some astrocytes (NSCs) of the adult brain readily give rise to neurons, while other astrocytes do not, despite expressing largely the same genes. We propose that striatal astrocytes do not usually give rise to neurons because their methylome locks them in their astrocyte fate by stabilizing the expression of astrocyte genes and by silencing neurogenic genes. On the contrary, vSVZ astrocytes receive signals present within the vSVZ that unlock their neurogenic fate and enable transition into qNSC2. NSCs in the vSVZ, once activated, have the epigenetic license to differentiate into neuroblasts, since many neurogenic genes are demethylated and thus more readily transcribed. Notably, injury elicits signals that unlock this potential outside the neurogenic niche. Yet, what makes some astrocytes become reactive, and others neurogenic, remains open. Likewise, it remains to be seen whether changes in DNA methylation are a driving force behind differentiation or whether the methylome is merely a stabilizing constraint on a particular differentiated fate. Targeted manipulation of DNA methylation, for instance using recently developed dCas9-based epigenome modifiers (Nakamura et al., 2021; Nuñez et al., 2021), may be employed to answer this question. Related to this, is the current strategy of reprogramming through forced expression of TFs similarly inducing remodeling of the methylome, or are there missing factors that would enhance the efficiency of reprogramming?

It is also currently unclear by which exact molecular mechanism methylation changes in the NSC lineage are introduced. Unexpectedly, we found little evidence for active demethylation via TET enzymes in the NSC lineage (Fig. 5d). One possible explanation is that our sample simply contained very few cells that were currently transitioning from vSVZ astrocyte to qNSC2, or that other mechanisms are involved. It is known that some TFs such as FOXA2 can trigger targeted demethylation (Donaghey et al., 2018; Reizel et al., 2021). It is thus possible that the TFs identified in Figure 2e have similar capabilities. Indeed, (Tchieu et al., 2019) recently demonstrated that one of our top hits enriched in astrocyte LMRs, NFIA, can be used to convert stem cells to astrocytes *in vitro*. The authors also report that this conversion involves demethylation at Gfap, a prominent astrocyte marker.

Intriguingly, the most significant methylation change that we observed after ischemia corresponds to a region in the Dnmt3a gene, which encodes the *de novo* DNA methyltransferase (Supplementary Fig. 5d-f). Given the surprisingly short timespan in which ischemia-induced methylation changes are reverted (Fig. 4c), demethylation at Dnmt3a might be part of a negative feedback loop in which ischemia first induces methylation changes at various loci including Dnmt3a. This in turn increases the expression of Dnmt3a, which then reverts the ischemia-induced demethylation events. Further research is required to investigate this hypothesis.

We propose that DNA methylation contributes to astrocyte cell identity. Surprisingly, we found that striatal astrocytes release this potential fate lock upon ischemic injury. This finding highlights the potential of endogenous astrocytes as therapeutic targets for regenerative medicine. Since the neurogenic capabilities of astrocytes in other brain regions are more limited (Magnusson et al., 2020), we hypothesize that the vSVZ, and to some extent the striatum, offers molecular cues required for changes in DNA methylation, as opposed to other brain areas. Furthermore, we showed that ischemia also causes a shift from vSVZ astrocytes towards qNSC2. Mathematical modeling of NSC dynamics led others to propose the existence of a resilient NSC population that does not usually contribute to neurogenesis (Harris et al., 2021; Ziebell et al., 2018). We hypothesize that this population is comprised within the vSVZ astrocytes and that their resilience is mediated by their methylome, while qNSC2 cells represent the homeostatic NSC pool. Along this line, we speculate that cues present during homeostasis or injury activate transition of distinct subsets of astrocytes into a primed qNSC2 state.

Finally, given the substantial number of epigenetic changes that occur upon ischemia, we wonder whether this innate astrocyte-to-neuron differentiation bears resemblance to *in vitro* reprogramming experiments such as astrocyte-to-neuron (Sharif et al., 2021) or iPSC-to-neuron conversion (Karumbayaram et al., 2009; Lindhout et al., 2020) via TFs. If so, it may be beneficial for reprogramming experiments to consider addressing the methylome to achieve stable and precise cell fates.

Our work shows that astrocytes acquire stem cell function through changes in DNA methylation. These changes occur in neurogenic areas of the adult brain, but can also be triggered by acute injury in common astrocytes. We also show that myelination is likewise regulated by changes in DNA methylation, many of which are dysregulated in MS patients. Targeting DNA methylation for endogenous cell replacement may be explored as therapeutic strategy to repair the diseased nervous system.

## STAR Methods

### Animals

In this work, the mouse lines C57BL/6N, and TiCY [B6-Tg(Nr2e1-Cre/ERT2)1Gsc Gt(ROSA)26Sortm1(EYFP)CosFastm1Cgn/Amv] (Liu et al., 2008) were used. All mice were male and were age-matched to 2 months old, except for the “ischemia 3 weeks” mice, which were three months old. Animals were housed in the animal facilities of the German Cancer Research Center (DKFZ) at a 12 h dark/light cycle with free access to food and water. All animal experiments were performed in accordance with the institutional guidelines of the DKFZ and were approved by the Regierungspräsidium Karlsruhe, Germany.

### TAM injection and ischemia

Two months old TiCY mice were intraperitoneally injected with tamoxifen (TAM). In these mice, tamoxifen-induced Cre recombination takes place in neural stem cells in the vSVZ, which express Tlx (Nr2e1) (Liu et al., 2008), and will stably activate the production of enhanced YFP, labeling NSCs and their progeny. Tamoxifen injection was done as described before (Carvajal-Ibañez et al., 2022). Two weeks after injection, an injury by bilateral carotid artery occlusion (BCCAO) was performed as described (Llorens-Bobadilla et al., 2015). The animals were sacrificed for single cell sorting either 48 h or 3 weeks after the ischemia.

### Single-cell suspension and FACS

For the ischemia experiment, the vSVZ and striatum were isolated. For the naïve experiments, vSVZ, striatum, and olfactory bulb were isolated. Depending on the plate, individual or pooled mice were used to sort cells on plate. For more information see Supplementary Table 2. Tissues were processed as described previously (Kremer et al., 2021) and sorted in a BD FACSAria II at the DKFZ Flow Cytometry Facility. Cells were stained with the following antibodies (all conditions and tissues together): O4-APC and O4-APC-Vio770 (Miltenyi; diluted 1:50), Ter119-APC-Cy7 (Biolegend; 1:100), CD45-APC-Cy7 (BD; 1:200), GLAST (ACSA-1)-PE (Miltenyi: 1:20), PSA-NCAM-PE-Vio770 (Miltenyi; 1:75), Prominin1-A488 (eBioscience; 1:75), and Sytox Blue (Life Technologies, 1:1000), CD9-eFluo450 (eBioscience, 1:300).

For sorting, we size-selected the vSVZ, striatum, or olfactory bulb cells and excluded for doublets, dead cells and CD45^+^/Ter119^+^ cells as recently described (Kalamakis et al., 2019). We then sorted different cell populations according to the tissue and experimental condition as follows: For naïve conditions and ischemic conditions: in the vSVZ we sorted Glast^+^ cells and O4^+^ cells, in the striatum we sorted Glast^+^ cells. Only for naïve conditions: In the olfactory bulb we sorted PSA-NCAM low and high cells. For the ischemia experiment, we additionally recorded the YFP information by performing index sorting. All the cells were sorted into individual wells of a 384-well plate. On early experiments we also sorted Glast/Prom1^+^ neural stem cells and O4^+^ oligodendrocytes.

### Miniaturized scNMT-seq protocol

For profiling the transcriptome and epigenome of single cells we developed and implemented a miniaturized and higher throughput version of the scNMT-seq protocol (Clark et al., 2018). On this new version, the Smart-seq3 (Hagemann-Jensen et al., 2020) method and specific normalization steps were implemented. A detailed version of the protocol is described in Cerrizuela et al., (2022).

### Methylase-assisted bisulfite sequencing (MAB-seq)

For performing MAB-seq, NSCs/astrocytes, oligodendrocytes and neuroblasts were sorted as follows: NSCs/astrocytes: Glast^+^/CD9^high^, oligodendrocytes: O4^+^, neuroblasts (PSA-NCAM^+^). For every biological replicate, 12 mice were pooled. Between 10,000 and 20,000 cells were sorted and processed following the MAB-seq protocol described in Wu et al. (2014).

### Processing of single-cell transcriptomic data

Transcriptomic reads were mapped to the mouse genome build GRCm38 (mm10) with STAR 2.7.3a (Dobin et al., 2013), using gene annotations downloaded from Ensembl Release 102. Both mapping and gene quantification were executed by the zUMIs pipeline 2.9.4f (Parekh et al., 2018) as described in the Smart-seq3 protocol (https://dx.doi.org/10.17504/protocols.io.bcq4ivyw).

### Processing of single-cell epigenomic data

Genomic reads were first trimmed with Trim Galore (https://www.bioinformatics.babraham.ac.uk/projects/trim_galore/) in paired-end mode, and then mapped to GRCm38 with Bismark 0.22.3 (Krueger & Andrews, 2011) in single-end, non-directional mode. After filtering PCR duplicates with Bismark, single-end alignments were merged. DNA methylation at individual cytosines was quantified with Bismark’s coverage2cytosine script using the NOMe-seq option to distinguish between CpG and GpC contexts.

### Quality filtering and analysis of single-cell epigenomic data

CpG methylation and chromatin accessibility (GpC methylation) was analyzed with `scbs` 0.3.2, a command line tool enabling the analysis of single-cell methylation data that was developed in parallel to this study. For a detailed explanation of the statistical methods, see (Kremer et al., 2022). In brief, we used `scbs prepare` to separately store CpG and GpC data in an efficient format and to compute quality metrics. Cells with read coverage of less than 50,000 CpG sites, or with poor methylation or accessibility profiles around their TSS were discarded. Methylation and accessibility profiles around TSSs and CTCF binding sites were computed with `scbs profile`. We furthermore discarded all cells with less than 2,500 observed genes. After filtering, we used `scbs smooth` with a bandwidth of 1000 (500 for GpC data) to quantify the smoothed mean methylation of all high-quality cells over the whole genome. VMRs and VARs were detected with `scbs scan`, a sliding window approach that scans the whole genome for regions of high methylation variance between cells. We used a bandwidth of 2000 (1000 for GpC data), a step size of 10 and a variance threshold of 0.2. We then quantified methylation and accessibility at VMRs, VARs and promoters (TSS ±1000 bp) using `scbs matrix`.

### Dimensionality reduction and pseudotime

We used Seurat 4.1.0 (Hao et al., 2021) to process the scRNA-seq data. To achieve higher resolution, we integrated our transcriptomic data with a much larger scRNA-seq dataset (wild-type cells from Carvajal-Ibañez et al. (2022)) as previously suggested (Argelaguet et al., 2019). Specifically, after normalizing and finding 3000 highly variable genes using default Seurat parameters for both data sets, we used FindIntegrationAnchors and IntegrateData using 30 dimensions to integrate the data sets, followed by scaling, PCA and UMAP on 30 PCs. We repeated the same approach to integrate the full data set containing cells from ischemic and naïve mice.

To visualize single cell methylomes, we subjected scaled and centered VMR methylation values to PCA, followed by UMAP on the top 15 PCs, excluding PC 5 which captured cell quality. Since epigenomic data contains missing values, we used a modified PCA that estimates missing values in an iterative manner (Kremer et al., 2022). To reduce noise, we used the “shrunken mean of residuals” reported by `scbs matrix` as a measure of methylation, and to reduce technical variation among cells we centered all values for each cell. Only VMRs observed in at least 20% of cells were used for PCA. Accessibility data was processed in the same manner, with the following differences: Only VARs observed in at least 40% of cells were considered, and promoter accessibility values were used in addition to VAR accessibility.

To project information from all three molecular layers into a shared lower-dimensional space, we used MOFA+ (Argelaguet et al., 2020) aiming for 15 dimensions (factors). As input, we used the expression values of 3000 highly variable genes (normalized with SCTransform (Hafemeister & Satija, 2019)), as well as the same methylation and accessibility values previously used for PCA (promoter methylation and accessibility, VMR methylation, VAR accessibility). We used UMAP on the top 13 MOFA factors, excluding factors 4 and 11 which captured technical variation. We then used Leiden clustering (Traag et al., 2019) on the MOFA factors, followed by slingshot (Street et al., 2018) to obtain pseudotime values informed by both gene expression and epigenetics.

### Correlation of epigenetic features with gene expression

Promoter methylation (shrunken residuals) and log-normalized gene expression values were correlated and tested for significance with the R function cor.test, using Pearson correlation. VMR methylation was correlated with the expression of the closest gene, as determined with bedtools (Quinlan & Hall, 2010): `bedtools closest -D` ‘`b`’ `-a regions.bed -b gene_bodies.bed`. Accessibility was correlated in the same manner. Only regions with genomic reads in at least 5 cells were considered. Correlation p-values were adjusted for multiple testing with the Benjamini-Hochberg method.

ChIPseeker (Yu et al., 2015) was used to quantify the number of VMRs and VARs that overlap with gene features, using the options tssRegion=c(−1000, 1000) and overlap=“all”. Overlaps with candidate cis-Regulatory Elements (ccREs, Registry V3, downloaded from https://screen.encodeproject.org/ on 2021-08-03, (Moore et al., 2020)) were quantified with the mergeByOverlaps function of the GenomicRanges R package (Lawrence et al., 2013).

### Quantifying methylation change along pseudotime

Correlation heatmaps of all three molecular layers were generated by binning cells along pseudotime, with a mean of 10 cells per bin. For all binned heatmaps, we enforced that each bin only contains cells from one cluster and tissue, so that e.g., the first cluster contains only striatal astrocytes. Methylation, accessibility, and expression values were averaged per bin and the Pearson correlation of all bins was visualized with ComplexHeatmap (Z. Gu et al., 2016).

To quantify (de)methylation events in the NSC lineage, we considered all VMRs that were observed in at least 100 cells of the NSC lineage. For each VMR, we fit a step function to the methylation values as a function of pseudotime. The function is parametrized by a change point *s* in pseudotime and two constant values, which the function takes before and after *s*. Minimizing the sum of squared residuals over this parameter space, we found a most likely value for the methylation change point in pseudotime. VMR change points were considered (de)methylation events if the step function fit was at least 15% better (with respect to the squared residuals sum) than a constant fit without a step.

### Epigenetic changes near cell-type specific genes

Representative marker genes for each cell type or stage were determined with the Wilcoxon rank sum test, by testing log-normalized expression values in cells of interest against the expression values of all other cells. We selected the top 100 most differentially expressed genes among genes with a Benjamini-Hochberg-adjusted p-value below 0.05 that also contain a VMR in their gene body. Expression/methylation/accessibility values of these genes and their corresponding promoters or VMRs were averaged.

### Transcription factor motif enrichment

Since the PCA on VMR methylation values captured methylation differences between oligodendrocytes and the neurogenic lineage on the first principal component (PC1), we selected the 5000 VMRs with the highest PC1 loading for oligodendrocyte-specific motif enrichment. We used HOMER (Heinz et al., 2010) with the JASPAR2022 motif database (Castro-Mondragon et al., 2022) to identify motifs enriched in these VMRs:

~~~
findMotifsGenome.pl VMRs.bed mm10r output/ -len 5,6,7,8,9,10,11,12 –size
given -mcheck JASPAR.db -mknown JASPAR.db -bg JASPAR.db
~~~

The same strategy was used to identify motifs enriched in regions with low methylation in the neurogenic lineage (5000 VMRs with the highest PC2 loading) and in astrocytes/vSVZ astrocytes (lowest 5000 PC2 loadings).

### Identification of lowly methylated regions (LMRs) and associated GO-terms

To identify regions that are differentially methylated between two sets of cells, we compared VMRs with the Wilcoxon rank sum test. Only VMRs with genomic coverage in at least 30 (5 when detecting ischemia LMRs, due to lower cell number) cells per cell set were considered. VMRs with an Benjamini-Hochberg-adjusted p-value below 0.05 were labeled LMRs. For visualizing gene expression in volcano plots and heatmaps, all LMRs overlapping a gene body were assigned to that gene. We used GREAT (McLean et al., 2010) for GO-term enrichment of genes near LMRs, using the option “basal plus extension” with a constitutive 0 kb upstream and 20 kb downstream regulatory domain, and up to 1000 kb max extension.

### Expression and methylation signatures induced by ischemia

Reactive astrocyte markers (Liddelow et al., 2017) were used to calculate expression signatures (mean log-normalized expression of the respective markers) for reactive astrocytes and the A1 and A2 subtypes. NSC-like methylomes and astrocyte-like methylomes were distinguished based on the mean methylation of all astrocyte and NSC LMRs; the methylation score represents the difference of these means.

### Processing and analysis of MAB-seq data

Reads were trimmed with Trim Galore 0.4.5 in paired-end mode and then mapped to GRCm38 with bwa-meth 0.2.2 (Pedersen et al., 2014) using default options. Reads with an unmapped mate, or with a mapping quality below 30, were discarded with samtools 1.9, followed by PCR duplicate removal with MarkDuplicates 2.18.14 from https://broadinstitute.github.io/picard/. Methylation values were extracted with MethylDackel 0.3.0 (https://github.com/dpryan79/MethylDackel) and common mouse single nucleotide polymorphisms were discarded with BS-SNPer (Gao et al., 2015) using the options –minhetfreq 0.1 --minhomfreq 0.85 --minquali 20 --mincov 10 --mapvalue 30.

To quantify active demethylation in LMRs that we had previously identified in our scNMT-seq data, we intersected the extracted MAB-seq methylation values with our LMRs using “intersect” from bedtools 2.27.1. All LMRs with a coverage greater than 5 in both replicates were then tested for evidence of active demethylation. To incorporate data from both replicates, we used a binomial generalized linear model (glm) with a logit-link function, testing the alternative hypothesis that the average methylation is lower than the average methylation level of the samples. p-values were adjusted for multiple testing with the Benjamini-Hochberg method.

MS DMRs from the supplementary material of Huynh et al. (2014) were converted from hg18 to mm9, using liftOver (https://genome.ucsc.edu/cgi-bin/hgLiftOver) with the option -minMatch=0.1, and then from mm9 to mm10 using default options. Converted MS DMRs were then glm-tested as described above.

## Author contributions

LPMK established a computational tool for analysis of sc-Methylation data and conducted computational analysis; SA supervised computational analysis; SC established high-throughput scNMT, generated scNMT profiles; MEAS and SC conducted global ischemia experiments; MEAS, TE, JS and AK supported establishment and conduction of multi-omics profiles; SD, SC, DW, and CP for generation of MAB-seq profiles; LPMK wrote the manuscript; SC, SA and AMV writing-review and editing; AMV conceived the research; LPMK, SC, SA and AMV formal design and conceptualization of the research; SA and AMV supervised and provided oversight of the project.

## Acknowledgments

We thank the DKFZ Genomics and Proteomics Core Facility, in particular, S. Wolf, the divisions of high-throughput sequencing, and the sequencing open lab; Jan-Philipp Malm from the DKFZ Single-cell Open Lab; Steffen Schmitt from the DKFZ Flow Cytometry Core Facility; Monika Langlotz from the ZMBH FACS Core Facility. This work was supported by the ERC (ERC-CoG 771376), the DFG (SFB873), the Klaus Tschira Foundation, the University of Heidelberg, and the DKFZ. We thank Andrés Sanz-Morejón for valuable feedback on the manuscript.

## List of Supplementary Materials

Supplementary Figures 1-5: embedded in the manuscript.

Supplementary Table 1: Metadata and quality metrics of 542 naïve cells and 614 ischemic cells.

Supplementary Table 2: Detailed information on all sequencing experiments.

Supplementary Table 3: List of marker genes used for the heatmaps shown in Supplementary Fig. 2b.

Supplementary Table 4: List of astrocyte LMRs, NSC LMRs, and oligodendrocyte LMRs.

